# Decontamination of Common Healthcare Facility Surfaces Contaminated with SARS-CoV-2 using Peracetic Acid Dry Fogging

**DOI:** 10.1101/2020.12.04.412585

**Authors:** Todd Cutts, Samantha Kasloff, David Safronetz, Jay Krishnan

**Author notes:** **Corresponding author:** Jay Krishnan, National Microbiology Laboratory, Public Health Agency of Canada, Winnipeg, Canada, R3E 3P6.

## Abstract

**Background:** The SARS-Cov-2 pandemic has highlighted the urgent need for safe and effective surface decontamination methods, particularly in healthcare settings.

**Methods:** The effectiveness of peracetic acid (PAA) dry fogging in decontaminating common healthcare setting surfaces was evaluated after experimentally contaminating nine surfaces (stainless steel, latex painted wood, unsealed hardwood, melamine countertop, vinyl flooring, clear plastic, faux leather, computer keyboard button and smartphone touch screen) with more than 10^6^ TCID_50_ of SARS-CoV-2.

**Results:** When fumigated with PAA dry fog for an hour, no infectious SARS-CoV-2 virus was recovered from experimentally inoculated coupons of representing nine different surface types. In contrast, high titer recovery of infectious virus was demonstrated for corresponding untreated drying controls of the same materials.

**Conclusion:** Standard surface decontaminating processes, including sprays and wipes, are laborious and often cannot completely decontaminate sensitive electronic equipment. The ease of use, low cost and overall effectiveness of a PAA dry fogging suggest it should be considered for decontaminating settings, particularly intensive care units where severely ill SARS-CoV-2 patients are cared for.

## Introduction

The burden of COVID-19 cases among healthcare workers has been staggering during the ongoing COVID-19 pandemic (1–5). Environmental sampling has demonstrated the presence of SARS-CoV-2, the causative agent of COVID-19, in indoor air and on various surfaces in healthcare settings (6–8). SARS-CoV-2 can persist on common surfaces for several weeks (9–11). Such contaminated surfaces could pose a significant risk of infection to healthcare workers and visitors (12). Surface decontamination using a variety of liquid disinfectants are routinely employed to disinfect various surfaces in healthcare facilities (13, 14). Disinfectants are generally applied as a spray or wipe which is labour intensive even on readily accessible surfaces and difficult, if not impossible, to apply on hard to reach surfaces. Employees who undertake liquid disinfectant application often are exposed to the hazardous chemicals in them (15, 16). Decontamination by fumigation using a gas, vapor, or fine mist is effective on all surfaces including the ones that are in the hard to reach areas; in addition, fumigation also decontaminates the air in the room. The objective of this study was to validate the efficacy of peracetic acid (PAA) dry fogging fumigation in decontaminating two rooms, and a variety of SARS-CoV-2 contaminated surfaces placed in them. Here we report the successful decontamination of two rooms and nine healthcare facility surfaces experimentally contaminated with SARS-CoV-2 using PAA dry fogging.

## Methods and materials

### Peracetic Acid Disinfectant

Liquid PAA is a strong oxidant and an excellent microbicide, its microbicidal capability has been known for more than a century (17); it can inactivate bacterial spores, fungi, and viruses (18–20). It is widely used in the food production/processing industry (21–23) because of its lack of toxic by-products. In the healthcare field, It has been used to disinfect endoscopes (24), sterilize bone allograft (25), and decontaminate surfaces in healthcare settings to control nosocomial infections, especially the ones caused by spore forming bacteria (26, 27). A number of PAA formulations have been registered with US Environmental Protection Agency and Health Canada as general disinfectants and also COVID-19 specific disinfectants (28, 29)

In 1968, PAA in vapor form was used to inactivate bacterial spores (30); in 2001 a fogger that created fine PAA particles smaller than 10μ was used for decontaminating hospital rooms and operation theaters (31). Unlike regular spray, the ultrafine particles of a fog do not readily settle on surfaces and cause dampness, hence the name dry fog. The dry fog behaves like a vapor, it fills the entire space, and diffuses into all areas and surfaces to provide decontamination of the whole space and the surfaces contained within. PAA fumigation has proven effective against bacterial spores and viruses (32–34), it has been used to decontaminate subway railcar (35), laboratories (33), biosafety cabinets (36), and N95 respirators (37, 38). In the air, the PAA has a half-life of 22 minutes (39), followed by breakdown to water, oxygen and carbon dioxide (39).

### Cell culture

African green monkey Vero E6 cells (ATCC CRL 1586; American Type Culture Collection, Manassas, VA, United States) were maintained at 37°C+5% CO_2_ in Cell Culture Medium (CCM) consisting of Dulbecco’s modified Eagle cell culture medium (DMEM; Hyclone SH3024302) supplemented with 10% Fetal Bovine serum (FBS; Gibco 12484028) and 1% v/v Penicillin /Streptomycin (PS, Gibco 10378016). Medium for virus cultures (VCM) consisted of DMEM supplemented with 2% FBS and 10 units per ml of PS.

### Stock virus preparation

Low passage SARS CoV-2 (hCoV-19/Canada/ON-VIDO-01/2020, GISAID accession# EPI_ISL_425177, kindly provided by the Vaccine and Infectious Disease Organization, VIDO, Saskatoon, Saskatchewan) was used to prepare concentrated stocks by infecting T-175 flasks of confluent Vero E6 cells at 0.01 multiplicity of infection. The health of the cell monolayer of the infected flask was compared to a noninfected Vero E6 flask over the course of the incubation. On day 3-4, cytopathic effect (CPE), as defined by cell detachment and cell rounding, became evident where over 90% of the cell monolayer was lifted in infected flasks. At this point, the supernatant was aspirated and pooled with a clarification step at low speed centrifugation (4500 g) for 10 minutes. The clarified supernatant was overlaid onto a 20% (w/v) sucrose cushion in Tris-NaCl-EDTA buffer and centrifuged at 134,000 g for 2 hours (Beckman Coulter 30 Ti rotor). The resulting viral pellet was suspended in VCM by repeat pipetting and aliquots stored in cryovials at minus 70°C until needed. All experimental work was conducted in high containment laboratories at the Canadian Science Centre for Human and Animal Health.

### Preparation of coupons

Nine surfaces that are commonly found in the healthcare settings were identified and used for this study: stainless steel, latex painted wood, unsealed hardwood flooring, melamine countertop, vinyl flooring, clear plastic, faux leather, computer keyboard button and smartphone touch screen. Small coupons (1-2 cm^2^) were cut where possible, while individual buttons were removed from an old computer keyboard. Three blackberry smartphones with touch sensitive screens were used to represent the omnipresent touch sensitive screens in healthcare facilities. Prior to use, all coupons were sterilized using gamma irradiation (1 Mrad, Cobalt-60 source), while the test surfaces of the smart phones were decontaminated using 70% ethanol wipes.

To prepare SARS-CoV-2 contaminated test surfaces, we followed an American Society for Testing and Materials (ASTM) International standard disinfectant testing method, ASTM E2197 (40). High titer SARS-CoV-2 virus (~ 5 × 10^8^ TCID_50_/mL) was mixed in a tripartite soil load (41) to create the test virus inoculum. The tripartite matrix which consisted of BSA, tryptone and mucin, represents the organic soil load: secretions/excretions within which the virus is released from an infected person. The inoculum was prepared fresh for each test replicate performed. Using a positive displacement pipette, 10 μl of inoculum was deposited onto the coupon surfaces and air-dried for 45-60 min in a biological safety cabinet.

### Dry fog fumigation assay

Fumigation experiments were carried out in a 164 m^3^ sized animal cubicle in a high containment laboratory; the cubicle consisted of two rooms with a door in between. The dry fogging system used for this study was described elsewhere (33); briefly, a portable dry fogger equipped with three AKIMist^®^ E nozzles that produces 7.5 μ sized droplets (Ikeuchi USA, Inc., Blue Ash, OH) was used. An air compressor (model 2807CE72; Thomas, Monroe, LA) set at 40 psi provided compressed air needed for the nozzles to generate the dry fog from the chemical contained in the 19L reservoir. The chemical, Minncare Cold Sterilant (Mar Cor Purification, Skippack, PA) had 4.5% PAA, which was diluted appropriately to achieve 1.6ml/m^3^ at 80% relative humidity (RH). Initial temperature and relative humidity levels of the rooms were measured using Professional Thermo-Hygrometer (TFA Dostmann Product#30.3039) and were used to calculate the amount of chemical and deionized water needed to be mixed to attain 80% RH. The door between the rooms was left open during the fumigation; the fogger was placed in the doorway with one nozzle directed towards the small room and the other two towards the large room (figure 1). Fourteen biological indicators (Spordex, #NA333, Steris, Mentor, OH) were placed at various locations within the rooms to validate room decontamination. Each biological indicator (BI) contained more than 10^6^ spores of *Geobacillus stearothermophilus* bacteria.

**Figure 1:**
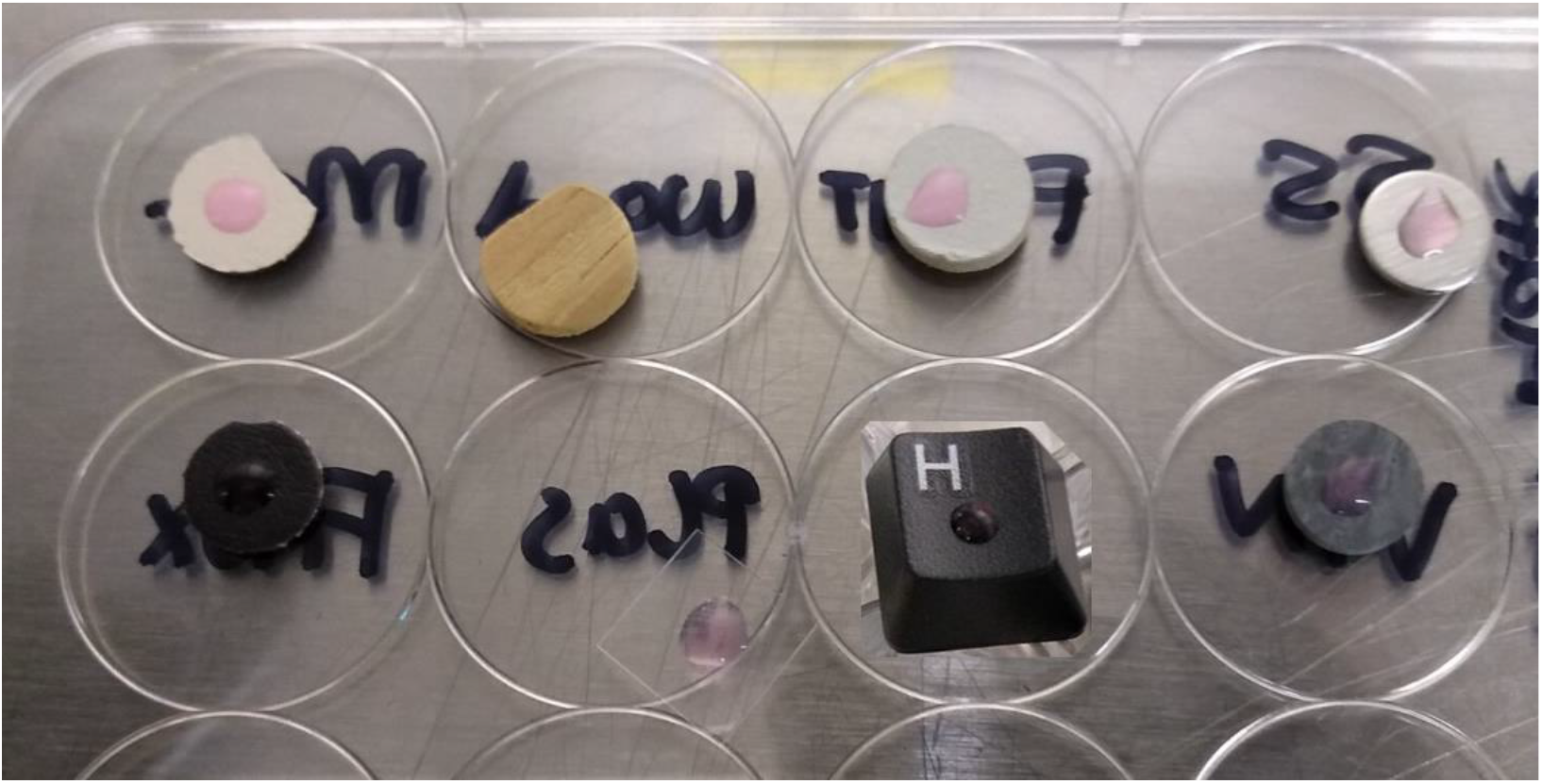
Preparation of SARS-CoV-2 contaminated test coupons from eight commonly found materials in healthcare settings. Coupons were inoculated with 10μl of SARS-CoV-2 virus (>10^6^ TCID_50_/coupon) mixed with a standard organic soil load and then allowed to dry in a BSC before being exposed to the PAA dry fog. Note the inoculum deposited on unsealed wood coupon got instantly absorbed. Top row (L to R): melamine, unsealed hardwood, latex painted particleboard, stainless steel Bottom row (L to R): faux leather, clear plastic, keyboard, vinyl flooring

One from each group of SARS-CoV-2 inoculated coupons was placed on the lids of three 12-well plates with their inoculated sides up (figure 2). The lids were positioned at a height of approximately 4 ft above the floor at three different locations in the room (figure 1). An exact set of triplicate coupons for each surface serving as unexposed positive controls were left in the BSC for the duration of the fumigation.

**Figure 2:**
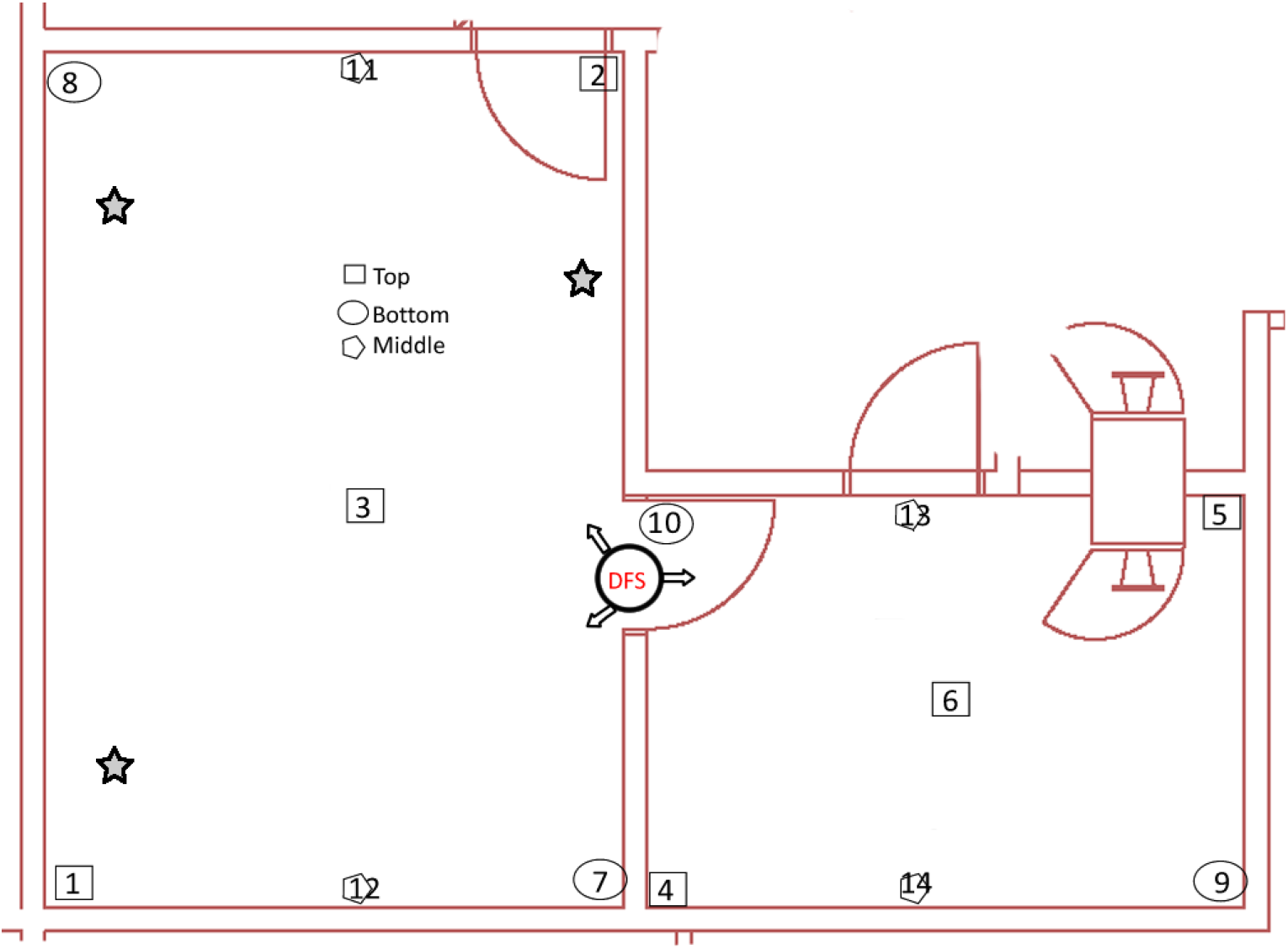
Preparation of the rooms for PAA dry fog fumigation. Stars indicate three different locations of SARS-CoV-2 contaminated test coupons, placed 4 ft above the floor. The DFS circle on the doorway between the rooms marks the location of the dry fogger; arrows indicate the directions of the fog nozzles. Numbers 1-14 indicate the locations of the biological indicators placed throughout the room to validate room decontamination.

Dry fogging process was initiated after turning off the laboratory air system, it took approximately 18-20 minutes and 2.5 litre of dilute chemical to reach 80% RH. After a contact time of one hour, the air system was turned on to aerate out the residual chemicals from the rooms; BIs and test surface coupons were retrieved for processing. BIs were incubated in trypticase soy broth at 56°C for 48 hours; an unexposed BI was also incubated similarly to serve as positive control for growth. The inoculum from each of the coupons was eluted in VCM along with their unexposed counterparts for quantification of viable virus titer in Vero E6 cells by TCID_50_. Three independent fumigation trials were carried out, each of which consisted of three replicates of each surface: three for fumigation and three as unexposed controls.

### Cytotoxicity control

As some coupon material could absorb residual PAA and/or contain chemicals from their manufacturing process, their potential negative impact on the cell monolayer (cytotoxicity) was also investigated. In replicates of three, each of the surfaces was exposed to the dry fog for 1 hour followed by 1-2 hr of aeration. Coupons were retrieved and subjected to the same elution protocol as test coupons. Eluates from each coupon was ten-fold serially diluted in VCM and added to Vero E6 monolayer in 96-well plates (50μl/well containing 150μl of VCM). Evidence of cytotoxicity to the cell monolayer was visually scored at day 5.

### TCID_50_ procedure

Vero E6 cells were seeded the day prior in a 96-well plate format to attain 80% confluence on the day of testing for virus titer by TCID_50_. Triplicate inoculated drying control coupons (unexposed control coupons) of each surface type as well as inoculated coupons that were exposed to PAA dry fog (exposed test coupons) were eluted into 1ml of VCM by repeat pipetting, each of which was then ten-fold serially diluted in VCM. Inoculated touchscreens were eluted in a total volume of 1ml of VCM by repeat washing of the inoculated area with 200ul VCM at a time. Media from the previously seeded 80% confluent Vero E6 cells were replaced with 150μl of fresh VCM prior to addition of the diluted virus inoculum. In replicates of 5 per dilution series, 50μl of diluted virus was added to Vero E6 cells and incubated at 37°C +5% CO_2_ for 5 days. Plates were examined for CPE under a light microscope and compared to a negative control to determine viral titer in TCID_50_ by the Reed Muench procedure (42).

## Results

Eluates obtained from fumigated clean coupons (cytotoxicity controls) showed no signs of cell death, except unsealed wood coupons (3/3 trials) and painted latex coupons (2/3 trials), which showed signs of cell death after overnight incubation when the undiluted eluates were added to Vero E6 cells. While titers of viable virus recovered from unexposed positive control coupons ranged between 10^4.5^−10^6.5^ TCID_50_/ml, no infectious virus was detected in tissue culture from any of the fumigated surface coupons in any of the three fumigation trials (figure 3). Triplicate coupons were tested in each independent fumigation experiment, all of which showed complete inactivation of SARS-CoV-2.

**Figure 3:**
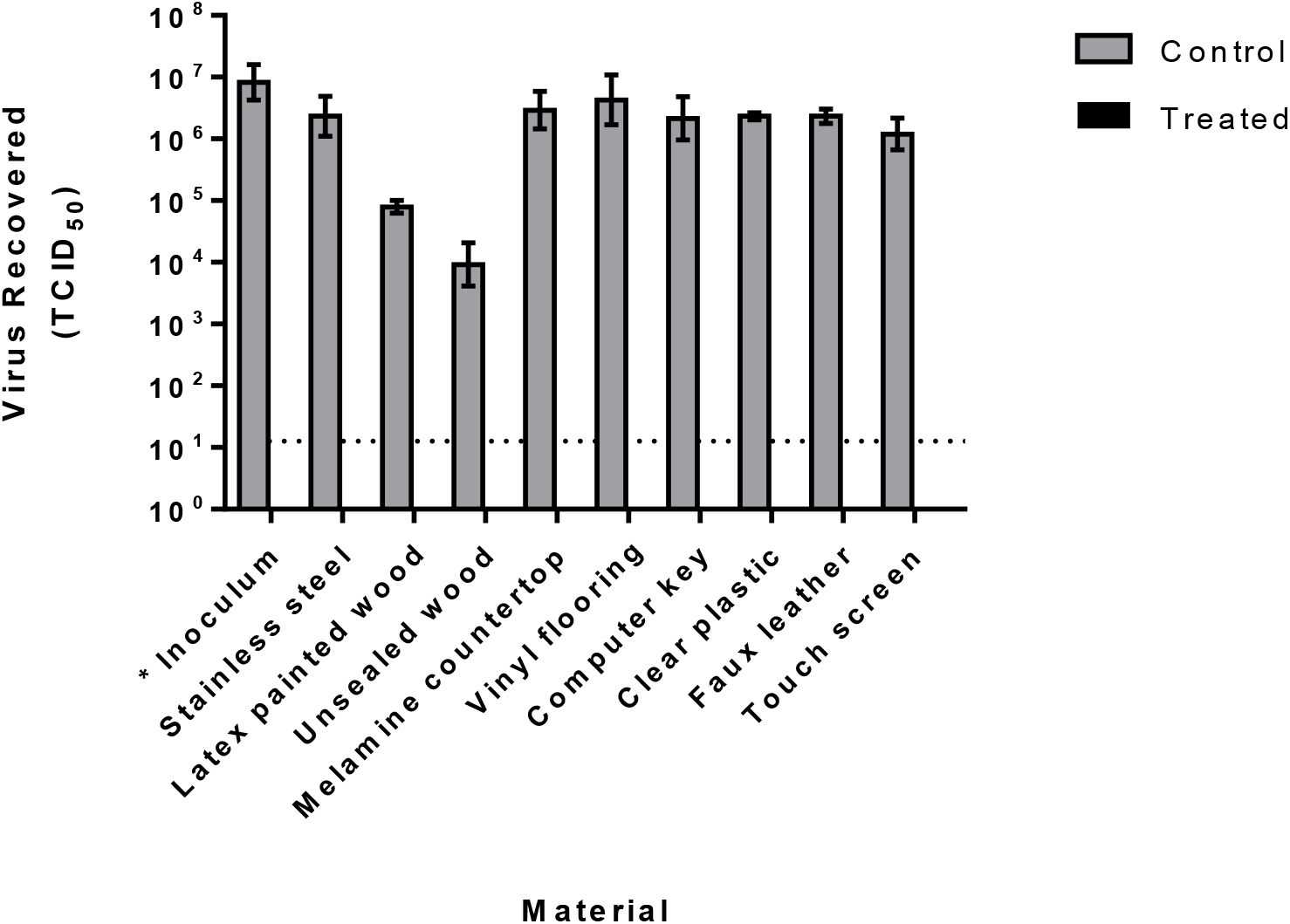
Inactivation of SARS-CoV-2 on nine common healthcare facility surfaces by peracetic acid dry fogging. Surface coupons contaminated with 10μl of virus inoculum were subjected to one hour dry fogging cycle (n=3 biological replicates per surface type) followed by elution in virus culture medium. No infectious virus was recovered from dry fog exposed coupons; viral titres recovered from the unexposed, dried positive control coupons of the same material type and quantified by end-point titration in Vero E6 cells are also shown. Dotted lines indicate limits of quantification for the TCID_50_ assay for unsealed wood and latex painted surfaces. Results represent means of three independent experiments.

For surface eluates which demonstrated cytotoxic effects to the Vero E6 cells (wood and painted latex), additional sub-passage of supernatants from the TCID50 plates was carried out to ensure that cell death observed at the neat dilutions of eluates from inoculated, PAA-treated surfaces was due to cytotoxicity rather than virus-induced CPE. Sub-passaging confirmed the lack of detectable infectious SARS-CoV-2 on both surface types in all experimental replicates. Interestingly, high titres of virus was recovered from unexposed control coupons made of non-porous materials while porous material coupons such as unsealed wood yielded lower concentrations (Figure 3), which is consistent with previous studies (43,44).

Developing the biological indicators failed to grow upon incubation for 48 hours, demonstrating that the entire two rooms were decontaminated along with the SARS-CoV-2 contaminated test surface coupons.

## Discussion

Widespread SARS-CoV-2 nosocomial infections have been reported from hospitals all across the world (45); according to WHO, healthcare workers accounted for 1 in 7 COVID-19 cases worldwide. SARS-CoV-2 transmission occurs via direct contact with infected persons, small airborne droplets, or larger respiratory droplets, or indirectly through contaminated surfaces/objects (fomite transmission). Heavily contaminated surfaces in environments housing infected patients present multiple sources of infection to healthcare personnel. One recent study (6) showed evidence of widespread contamination on surfaces in patient room including toilets, ventilation grills, and even on the floor under the beds without direct patient contact. Therefore, it is critical that the decontamination methods adopted should reach all surfaces in the room, including those in hard to reach areas.

Routine surface decontamination processes using liquid sprays/wipes is labor intensive, often hazardous to decontamination personnel, and can not reach all hard-to-reach surfaces. Fumigation on the other hand, keeps the personnel out of the room being fumigated while it decontaminates the entire room including the air and the various surfaces contained within. Infectious agents do not deposit themselves cleanly on the surfaces, they will be in a milieu of patients’ excretions/secretions, which after drying would be a challenge for disinfecting chemicals to inactivate. In this study, the SARS-CoV-2 virus was suspended in a standard tripartite organic soil load to represent such a challenging milieu, and then dried on to the test coupons. As noted before, PAA both in the liquid and the fumigant form (33, 46), tolerates organic soil load well, which is consistent with our finding here, where all the surfaces were decontaminated upon a one hour exposure. Expensive electronic equipment is plentiful in modern healthcare settings, they are a necessity to provide state of the art patient care. A fumigation technology selected to decontaminate such a facility and the equipment in it should not damage electronic equipment, PAA fumigation has previously been shown to be compatible with electronics after repeat exposures in a laboratory setting (33).

Dry fog fumigation using PAA is a low tech, cost effective and portable decontamination technology that can be used for decontaminating large areas within a short period. This study shows that PAA fumigation resulted in the complete inactivation of SARS-CoV-2 on all the nine test surfaces as well as the decontamination of the rooms that housed them. While the focus of this work was decontamination of surfaces found in healthcare settings, these materials are common in a variety of structures. There have been reports of COVID-19 outbreaks in cruise ships (14, 47), schools (48), sports facilities (49), and long-term care centres (50). Thus, PAA fumigation can be used to successfully decontaminate not only healthcare facilities, but also a variety of other indoor spaces and facilities.

